# Programming Targeted Immune Cell Migration Using Orthogonal Chemokine-Receptor Pairs

**DOI:** 10.64898/2026.10.12.758398

**Authors:** Jungmin Lee, Karthik Srinivasan, Claudia Llinas del Torrent, Debraj Ghose, Devan H. Shah, Luis Oliva Hernandez, Sarah Wasinger, Michelle J. Chang, Wei Yu, Yurie Tonai, Francis C. Peterson, Martyna Szpakowska, Andy Chevigné, Brian F. Volkman, Aashish Manglik, Wendell A. Lim

## Abstract

Chemokines orchestrate the migration of immune cells throughout the body, but harnessing chemokine signaling to engineer targeted cell trafficking is challenging because of their overlapping interactions and broad expression patterns. Here we used structure-guided protein design to engineer orthogonal chemokine-receptor pairs with minimal crosstalk to native systems. T cells engineered with an ortho-receptor selectively infiltrated tissues producing the ortho-chemokine, in vitro and in vivo. By engineering T cells that both sense and secrete the ortho-chemokine, we generated collective swarming behavior in which pioneer cells attract follower cells via an ortho-chemokine gradient self-organized by positive feedback. By providing orthogonal genetic control over both the generation and sensing of chemokine gradients, this system enables the construction of autonomous spatial targeting programs in engineered cells.

## Main Text

Chemokines guide the migration and spatial organization of immune cells in the body – they recruit neutrophils to sites of infection, direct effector T cells into inflamed tissues, and organize lymphocytes into specialized zones within lymphoid organs (*1, 2*). Complex and dynamic cell migration patterns are determined by what cells produce a chemokine, and what cells express the cognate chemokine receptor (which belong to the Gi-coupled GPCR family). Chemokine signaling is an attractive lever for engineering cell trafficking, if it could be exploited to spatially direct immune cell therapies to sites of disease. The endogenous chemokine network, however, is highly degenerate, with ∼50 chemokines and 23 receptors forming >100 functional interactions (*3*). Moreover, both chemokines and receptors are broadly expressed across multiple tissues (*4*). Thus, using native chemokine systems to predictably and selectively engineer cell trafficking is challenging. For example, using the chemokine CXCL9/10/11 to recruit effector T cells to tumors could also lead to recruitment of Tregs and myeloid-derived suppressor cells (*4–6*).

It would be much more powerful to have a synthetic orthogonal chemokine-receptor pair that could be used to engineer novel immune cell trafficking programs, without crosstalk and interference with the native chemokine network. Other synthetic receptors, such as chimeric antigen receptors (CARs) and synthetic Notch (synNotch) receptors, have proven remarkably effective at programming novel cellular responses (*7–9*). We hypothesized that a chemokine-receptor pair engineered to be mutually orthogonal to the native network could enable predictable, cell-autonomous control over both chemokine secretion and sensing, and thus, could be a foundational module for engineering new cell migration programs.

Here we used structure-based protein design to engineer orthogonal chemokine-receptor pairs that show minimal crosstalk with the native network. These signaling pairs could direct selective T cell migration and tumor infiltration, in vitro and in vivo. Moreover, we could use these components to engineer a self-amplifying, collective T cell swarming circuit that operates through a synthetic positive feedback loop in which cells both sense and secrete the ortho-chemokine. This orthogonal chemokine-receptor system is one of the first examples of an orthogonal GPCR-ligand pair that provides genetically encoded control of both signal production and sensing. Thus, this system establishes a platform for the construction of novel spatial targeting programs, with diverse potential applications in cellular therapies for cancer, autoimmunity, and beyond.

## RESULTS

### Orthogonalization of the CCL20/CCR6 chemokine/receptor interface via structure-guided charge swap mutagenesis

To design orthogonal chemokine-receptor pairs, we modified the binding interface of a native chemokine-receptor pair by introducing complementary mutations to both the receptor and the ligand (**Fig. 1A**). We started with the CCL20-CCR6 chemokine-receptor pair as a template, as it is one of the few monogamous signaling pairs and has a relatively small, shallow interface to target for orthogonalization. We reasoned that limited engagement of the receptor core by the chemokine would facilitate orthogonalization, avoiding redesign of the extensive buried hydrogen-bonding networks found in other chemokine–receptor complexes. Based on the cryo-electron microscopy (cryo-EM) (*10*) and AlphaFold2 structures, we identified a polar hydrogen bond network at the interface involving a single residue (E30) in the 30s loop of the chemokine interacting with two residues (T191, K200) in the extracellular loop 2 of the receptor (**Fig. 1B**). To engineer an orthogonal pair, we used a compensatory charge-swap strategy in which the polar interactions at the wildtype interface were simultaneously reversed at all three contact positions, preserving the overall binding geometry while redirecting specificity toward a matching mutant pair (**Fig. 1C**).

**Fig. 1.**
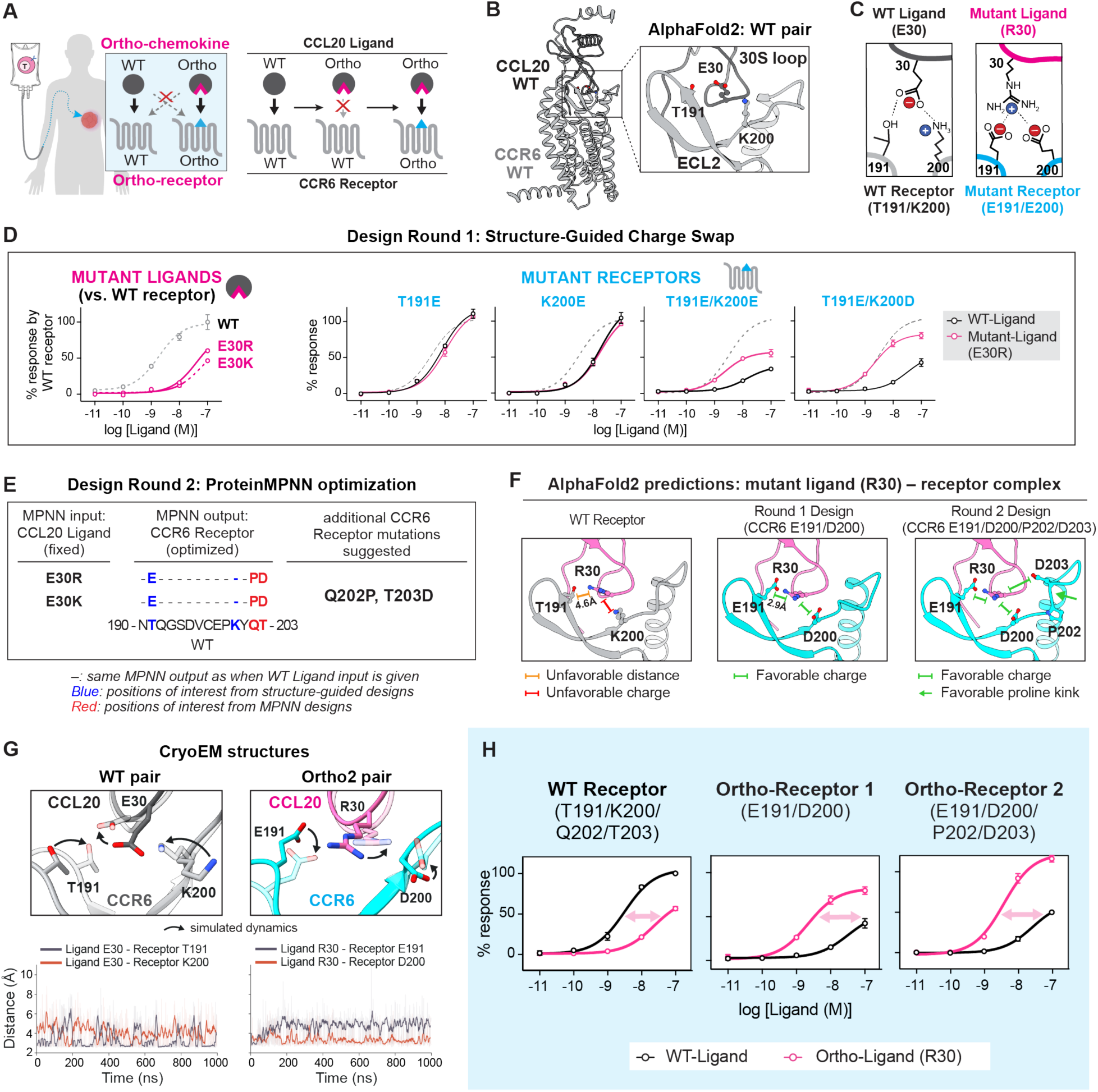
Structure-guided design of orthogonal chemokine-receptor pairs. (**A**) Design strategy: complementary mutations at the CCL20-CCR6 interface favor mutant-mutant interaction while disrupting binding to wildtype partners. (**B**) AlphaFold2 model of wildtype CCL20-CCR6 showing ligand E30 forming a hydrogen bond network with receptor T191 and K200. (**C**) Charge-swap mutations at ligand E30 and receptor T191/K200 invert local electrostatic complementarity to favor mutant-mutant over mutant-wildtype interactions. (**D**) Charge-swap mutant ligands and receptors show reduced signaling via their wildtype partners. Double mutant receptors (T191E/K200E, T191E/K200D) show selectivity for E30R ligand, with T191E/K200D receptor showing higher maximal efficacy (NanoBiT Gαi recruitment; N=2–3; Mean±S.E.M.). (**E**) ProteinMPNN identifies additional receptor mutations, Q202P and T203D. (**F**) AlphaFold2 models show MPNN-designed receptor mutations further improve charge and structural complementarity with mutant ligand. (**G**) Cryo-EM structures and associated simulations reveal stable CCL20-CCR6 engagement in both wildtype and mutant (“Ortho”) pairs. Side chain density at the interface was weak, so the indicated side chain conformations were modeled tentatively. Transparent models indicate alternate conformations sampled during MD simulations. Representative snapshots illustrate interactions between ligand E30 (R30 in Ortho) and receptor T191/K200 (E191/D200 in Ortho). Corresponding inter-residue distances over 1-μs MD simulations are shown below. (**H**) Ortho-Ligand (R30) preferentially activates Ortho-Receptors 1 (E191/D200) and 2 (E191/D200/P202/D203) while showing reduced signaling through the wildtype receptor (NanoBiT Gαi recruitment; N=2–3; Mean±S.E.M.).

Our first step was to functionally disrupt the chemokine ligand side of the interface. To achieve this goal, the ligand was engineered with a charge swap mutation (E30R or E30K). These mutant ligands showed 20–30-fold reduced activation of the wildtype receptor in a NanoBiT Gαi recruitment assay (**Fig. 1D**). As a second step, we sought to identify mutations on the receptor side of the interface that could rescue interaction with the mutant ligand. We designed potentially compensatory mutant receptors and assessed their ability to be activated by the E30R mutant ligand. Receptors were mutated at either or both of the T191 and K200 positions to a negatively charged amino acid, to compensate for the positively charged ligand mutation. Mutant receptors with a single mutation (T191E or K200E) were insufficient to discriminate the mutant ligand from the wildtype ligand (**Fig. 1D**). A double mutant receptor (T191E/K200E), however, was preferentially activated by the mutant ligand over the wildtype, although the maximal efficacy was reduced approximately two-fold compared to the wildtype pair (**Fig. 1D**). To identify a double mutant receptor with improved efficacy, we screened a set of mutant receptors with varying combinations of mutations at these two positions, testing different negatively charged residues or isosteric amide residues (**fig. S1**). We found that the T191E/K200D combination showed the best selectivity for the mutant ligand over the wildtype, while maintaining the maximal efficacy comparable to the wildtype pair (**Fig. 1D**).

### ProteinMPNN structure-guided optimization of orthogonal interface

To further optimize this orthogonal interface, we used ProteinMPNN (*11*) to survey additional mutations in the receptor extracellular loop 2. This approach identified two additional receptor mutations (Q202P, T203D) predicted to geometrically complement the ligand mutation (E30R or E30K), generating a quadruple mutant receptor (T191E/K200D/Q202P/T203D) (**Fig. 1E, F, fig. S2A**). ProteinMPNN also independently nominated the same ligand substitution (E30R and E30K) when predicted against several mutant receptors including the quadruple mutant receptor as structural input, matching our structure-guided design (**fig. S2B**).

Together, these results converged on our final designs and nomenclature: the E30R mutant ligand (“Ortho-Ligand”), the T191E/K200D double mutant receptor (“Ortho-Receptor 1”), and the T191E/K200D/Q202P/T203D quadruple mutant receptor (“Ortho-Receptor 2”). Ortho-Ligand preferentially activated Ortho-Receptor 1 and Ortho-Receptor 2, while showing reduced activation through the wildtype receptor (**Fig. 1H**).

### Structural comparison of WT and Orthogonal chemokine-receptor complex

We determined cryo-EM structures of the wildtype chemokine-receptor pair (3.9 Å resolution) and the engineered chemokine-receptor pair comprising the Ortho-Ligand and the Ortho-Receptor 2 (3.6 Å resolution) (**Fig. 1G, figs. S3, S4**). The two structures showed comparable ligand-bound active conformations (**figs. S5, S6**), indicating that mutagenesis in the ligand binding interface did not disturb the overall conformation essential for receptor activation and downstream signaling. Because side-chain density at the interface was weak, we performed molecular dynamics (MD) simulations on the wildtype chemokine-receptor pair and the orthogonal pair involving Ortho-Receptor 2 to explore possible side chain interactions at the engineered interface. These simulations suggested that the same three positions (ligand residue 30; receptor residues 191 and 200) mediate the interface in both pairs, but with reversed charge distribution, consistent with our design strategy. Interestingly, the two pairs showed distinct interaction dynamics: the wildtype interaction (E30-T191-K200) was structurally more flexible, with ligand residue E30 alternating contacts between receptor residues T191 and K200, whereas the corresponding mutant interaction (R30-E191-D200) was highly stable, with ligand R30 stably engaged with receptor residue D200 (**Fig. 1G**). In contrast, MD simulations of non-matching wildtype-ortho pairs showed highly fluctuating side-chain conformations and did not sustain close inter-residue contacts over the simulation timescale (**fig. S7**).

### Orthogonal chemokine system overcomes cross-reactivity of native chemokine network

The engineered Ortho-Ligand/Receptor pairs were selected for their ability to disfavor interaction with their parental wildtype partners, the closest relatives most likely to cause crosstalk. But we also wanted to confirm that these mutations did not lead to novel crosstalk – i.e. that the Ortho-Ligand did not induce novel cross-activation of any other native chemokine receptors; and that the Ortho-Receptor did not respond to any other native chemokines. To test whether the Ortho-Ligand/Receptor pairs are truly orthogonal to the entire human native chemokine network, we used NanoBiT β-arrestin1 recruitment assays to perform a complete screen for any native chemokines that could activate Ortho-Receptors and any native receptors that could respond to Ortho-Ligand (we also performed a parallel screen using the parental chemokine CCL20). We found that neither Ortho-Receptor was substantially activated by any other 44 native human chemokines, with Ortho-Receptor 2 showing the best orthogonal profile. Similarly, the Ortho-Ligand only strongly activated Ortho-Receptors 1 and 2, and did not significantly activate any other native signaling receptors (**Fig. 2, fig. S8A, S8C, S8E**). To assess whether Ortho-Ligand/Receptor pairs would be non-crossreactive in mouse models, we also tested their interaction with the murine “parental” counterparts (mCCL20/mCCR6) and found no interaction (**fig. S8B**).

**Fig. 2.**
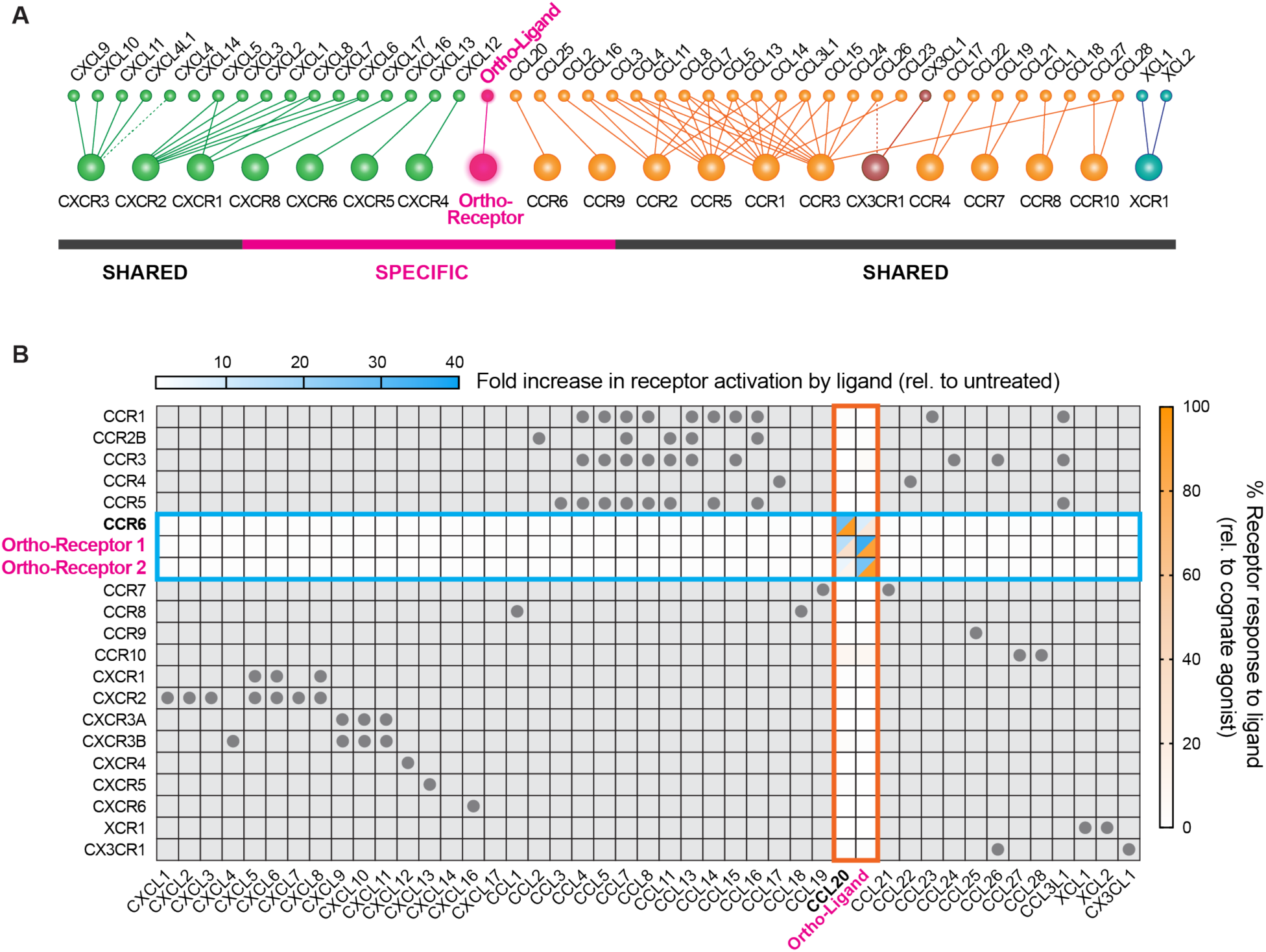
Engineered chemokine/receptor pairs are orthogonal to the native chemokine/receptor network with minimal crosstalk. (**A**) Chemokine network diagram highlighting the specific Ortho-Ligand/Ortho-Receptor interaction, in contrast to the shared interactions of native chemokine-receptor pairs. (**B**) Heat map of chemokine-receptor interactions. Ortho-Ligand and Ortho-Receptors 1 and 2 are specific for each other and do not interact with native components. Only pairs within the blue and orange boxes were experimentally tested in a NanoBiT β-arrestin1 recruitment assay using 100 nM chemokine. Chemokine screening data (x-axis, blue scale) were normalized to untreated controls; receptor screening data (y-axis, orange scale) were normalized to each receptor’s response to its cognate agonist chemokine (**table S1**). Dots indicate known interactions reported in the literature (*3, 12, 13*).

To further confirm that the Ortho-Ligand does not significantly attract any native chemotactic immune cells, we applied human peripheral blood mononuclear cells (PBMCs) to tumor spheroids secreting wildtype CCL20 or Ortho-Ligand (mutant CCL20). We found that endogenous immune cell types that migrated toward spheroids expressing the wildtype CCL20 chemokine (such as Tregs, B cells, and dendritic cells) were not recruited to spheroids expressing Ortho-Ligand (**fig. S9**). Interestingly, biochemical analysis showed that Ortho-Ligand still retained interaction with the atypical chemokine receptors (ACKRs) that interact with WT CCL20 (ACKR4 and ACKR5) (**fig. S8D, E**). The ACKRs are sink receptors that play a key role in shaping chemokine gradients (*12, 13*). Together, these results indicate that the Ortho-Ligand/Receptor pairs are orthogonal to the native signaling network, while Ortho-Ligand also retains sink receptor interactions that may preserve the native chemokine’s ability to form spatial gradients in vivo.

### Orthogonal chemokine system enables selective T cell trafficking to a spatially defined source

To test whether Ortho-Ligand/Receptor pairs can guide directed migration of primary T cells in a selective manner, we first used a gradient chamber slide (ibidi) to expose T cells to a chemokine concentration gradient in Matrigel, mimicking an extracellular matrix-like environment.

Engineered T cells showed chemokine-dependent directional migration. Ortho-Receptor-expressing T cells migrated only toward Ortho-Ligand and not WT-Ligand, and Ortho-Ligand attracted only Ortho-Receptor-expressing T cells and not WT-Receptor-expressing or untransduced T cells (No Receptor) (**Fig. 3A-B, Movie S1**). Quantification of the forward migration index further showed that T cells expressing Ortho-Receptor 1 exhibited a slightly stronger directional migration toward Ortho-Ligand than those expressing Ortho-Receptor 2 (**Fig. 3C**).

**Fig. 3.**
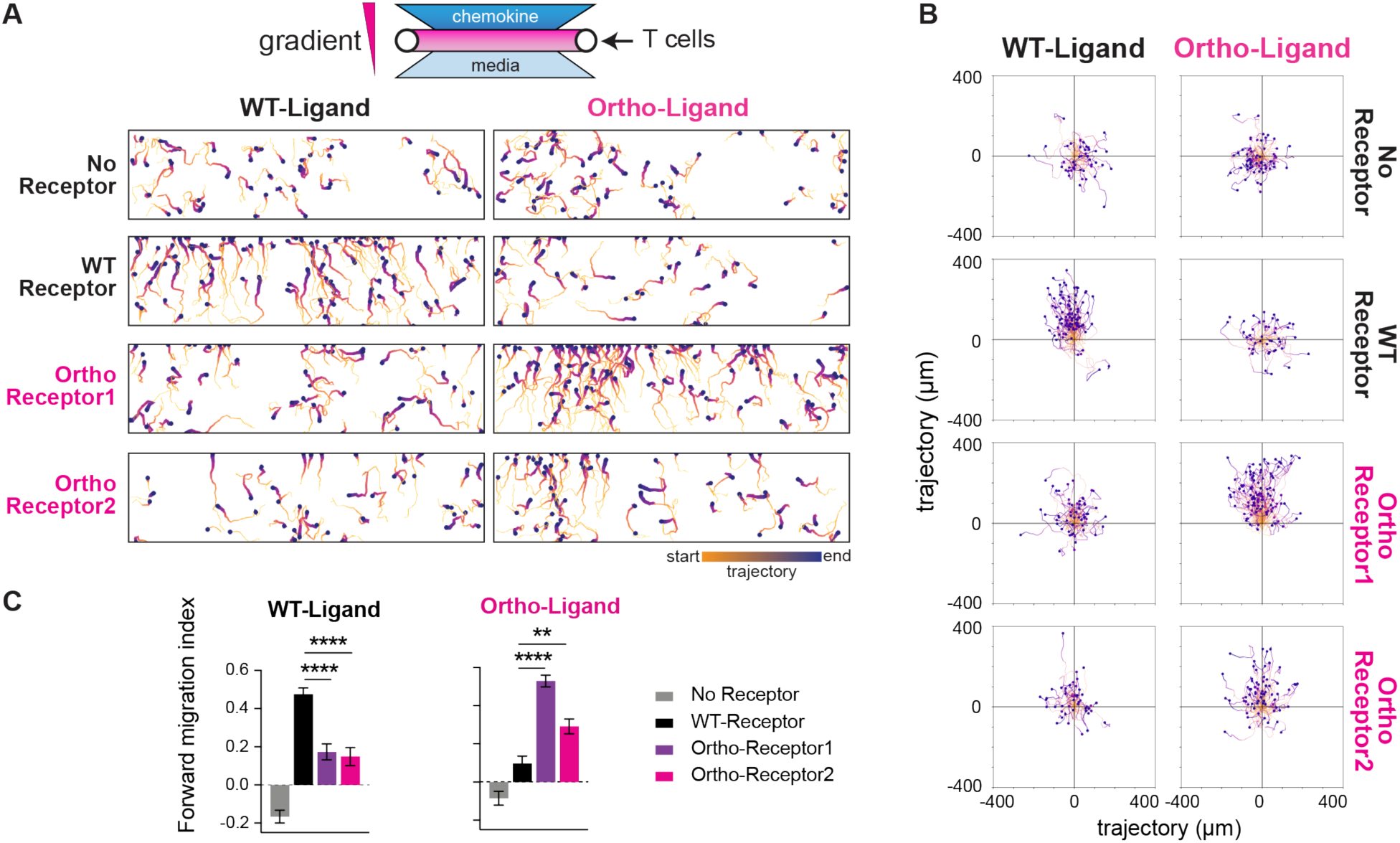
Ortho-Ligand/Receptor pairs induce directed migration of engineered T cells orthogonally to wildtype. (**A**) T cells expressing Ortho-Receptor 1 or 2 migrate toward Ortho-Ligand but not wildtype ligand in a μ-slide chemotaxis system (ibidi) over 1–1.5 hours. Each line shows the migratory track of a single cell in the upper half of the middle chamber. Increasing darkness and thickness indicate later timepoints. Only cells that moved at least 50 μm (5 cell diameters) are shown. 50 nM chemokine was applied to the upper gradient chamber. (**B**) Trajectory plots of individual T cells from (A), normalized to the origin (0, 0). (**C**) Forward migration index shows efficient directed migration along the chemokine gradient axis (y-axis) only when T cells are exposed to the ligand matching their expressed receptor. N=84–150. Mean±S.E.M. One-way ANOVA test. **p<0.01, ****p<0.0001.

Next, we asked whether Ortho-Ligand/Receptor interactions confer spatial selectivity for T cell migration across distinct spatial compartments. We tested this using a spheroid infiltration assay, in which T cells were seeded in the Matrigel surrounding an A549 tumor spheroid which was engineered to secrete WT-Ligand or Ortho-Ligand (**fig. S10A**). A similar spheroid not expressing any chemokine was used as a control. When the number of chemokine producer cells in each spheroid was systematically titrated, engineered T cells infiltrated into spheroids in a dose-dependent manner, showing the selectivity and chemotactic strength of each receptor for WT- or Otho-Ligand consistent with the previous results (**fig. S11**).

When all three spheroids – a control spheroid, WT-Ligand-producing one, and Ortho-Ligand-producing one – were placed in the same well, creating two spatially distinct chemokine gradients, engineered T cells infiltrated preferentially into the spheroid secreting the cognate chemokine (recognized by their expressed receptor). Notably, Ortho-Receptors 1 and 2 showed differing responses to WT-and Ortho-Ligand. T cells expressing Ortho-Receptor 1 infiltrated both spheroids, indicating higher cross-reactivity with WT-Ligand, whereas those expressing Ortho-Receptor 2 showed a stronger preference for the Ortho-Ligand-producing spheroid, indicating higher specificity (**Fig. 4A, Movie S2**).

**Fig. 4.**
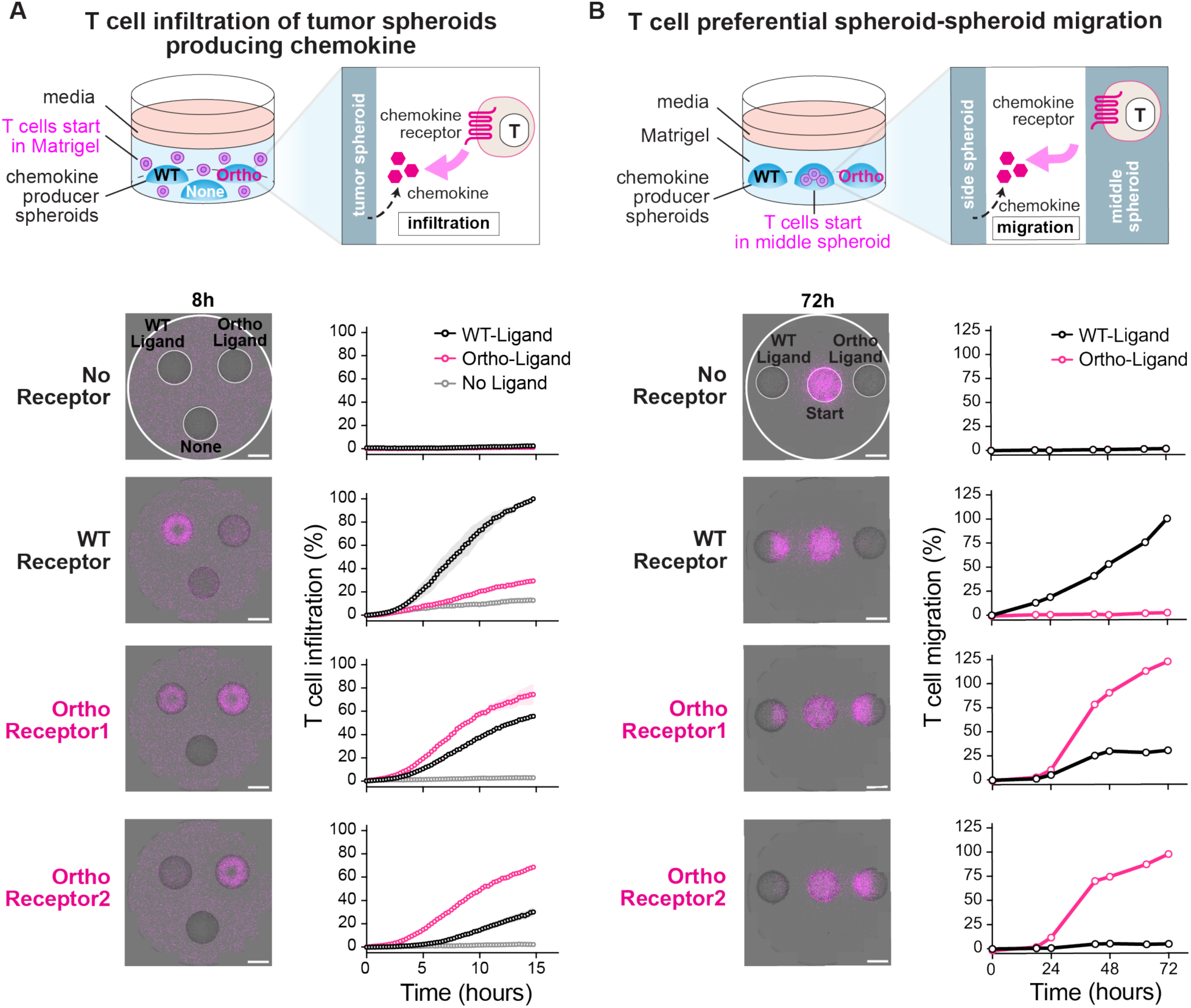
Ortho-Ligand competitively recruits engineered T cells to a spatially defined source in vitro. (**A**) Three A549 tumor spheroids – each secreting WT-, Ortho-, or no chemokine – are seeded in each well of a 24-well plate, along with T cells dispersed in an overlaying Matrigel layer. T cells infiltrate only into the spheroid secreting the chemokine cognate to their expressed receptor. Images taken 8 hours after setup. Scale bars, 2 mm. Representative of three independent experiments. N=2. Mean±S.E.M. See **Movie S2** for a movie of T cell infiltration into spheroids. (**B**) Three tumor spheroids are seeded linearly, with T cells starting in the middle spheroid (mixed with parental A549 cells) and WT-or Ortho-Ligand-secreting spheroids on either side. T cells egress from the middle spheroid and migrate toward the side spheroid secreting their cognate chemokine. Images taken at 72 hours. Scale bars, 2 mm. Representative of three independent experiments. See **Movie S3** for a movie of T cell migration between spheroids.

In an alternative three-spheroid competition arrangement, T cells were pre-embedded in a middle spheroid, surrounded by two other spheroids each secreting either WT- or Ortho-Ligand. Engineered T cells migrated toward one neighboring spheroid over the other in a chemokine-dependent manner, competitively selecting their cognate chemokine source (**Fig. 4B, Movie S3**). These studies again showed optimal preference of Ortho-Receptor 2 for the Ortho-Ligand over the WT-Ligand. In subsequent studies we focused only on Ortho-Receptor 2, as this receptor showed the most orthogonal behavior.

### Orthogonal chemokine system directs T cell trafficking and localization in vivo

In vivo, immune cells navigate complex tissue environments to traffic to target tissues, guided by spatially defined chemokine gradients. To test whether the orthogonal chemokine system can guide systemically administered T cells specifically to target sites in vivo, we implanted bilateral subcutaneous A549 tumors in mice, with one tumor engineered to express Ortho-Ligand and the contralateral control tumor engineered to express WT-Ligand or mCherry, all under doxycycline-inducible control (**fig. S10B**). To induce chemokine production in the tumors, mice were given doxycycline chow, and one day later received T cells expressing Ortho-Receptor 2 and an effLuc reporter via IV tail vein injection.

In these mouse experiments, Ortho-Receptor 2-expressing T cells preferentially accumulated in the Ortho-Ligand expressing tumor, whether the opposing tumor expressed WT chemokine (**Fig. 5A, fig. S13**) or no chemokine (mCherry) (**fig. S12**). Notably, Ortho-Receptor 1-expressing T cells were found to accumulate in both Ortho-Ligand and WT-Ligand expressing tumors, consistent with the earlier in vitro results indicating that this receptor shows responses to both ligands. These studies confirmed that the Ortho-Ligand/Receptor 2 pair can selectively guide T cell trafficking over long distances in vivo.

**Fig. 5.**
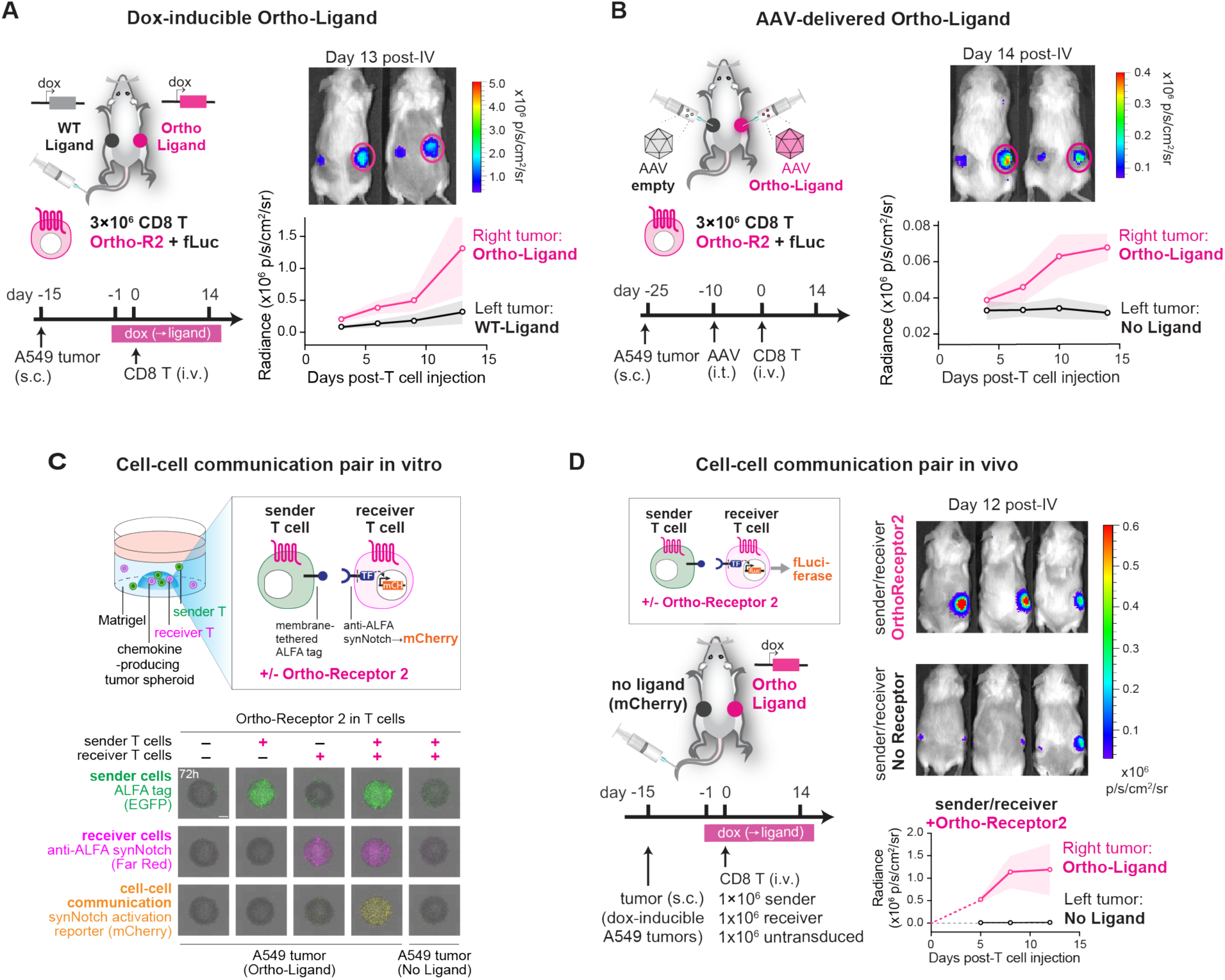
Ortho-chemokine system can guide T cell trafficking in vivo. (**A**) NSG mice bearing bilateral A549 tumors, each secreting WT-or Ortho-Ligand under doxycycline induction, received T cells via tail vein IV. Images from day 13 post-T cell injection. (**B**) Mice bearing bilateral parental A549 tumors are injected intratumorally with AAV encoding a mock payload or Ortho-Ligand prior to T cell IV injection. Images from day 14 post-T cell injection. In (A) and (B), T cells expressing Ortho-Receptor 2 preferentially traffic to the Ortho-Ligand-secreting tumor. T cells express effLuc and are tracked by bioluminescence imaging over 14 days post-T cell injection. N=4–5 mice per group. Mean±S.E.M. (**C**) In a 96-well plate, A549 spheroids secreting Ortho-Ligand are overlaid with Matrigel containing two T cell populations: Sender T cells (membrane-tethered ALFA tag) and Receiver T cells (anti-ALFA synNotch family receptor, specifically SNIPR (*13*), driving mCherry expression). Sender (green) and receiver (magenta) cells activate SNIPR (mCherry, yellow) only when both express Ortho-Receptor 2. Images are from 72 hours. Scale bar, 1 mm. (**D**) Mixed Sender/Receiver populations, with or without Ortho-Receptor 2, are injected into mice bearing bilateral A549 tumors expressing mCherry or Ortho-Ligand under doxycycline induction. Both populations co-traffic to the Ortho-Ligand-secreting tumor only when both express Ortho-Receptor 2, driving SNIPR-induced effLuc expression. N=3–4 mice per group. Mean±S.E.M.

To explore the viability of alternative ortho-chemokine expression strategies, we replaced the engineered tumors with unmodified parental tumors and instead delivered the Ortho-Ligand gene locally via intratumorally injected adeno-associated virus (AAV). We confirmed that the transduced tumors secreted Ortho-Ligand (**fig. S14A, B**). Consistent with the previous experiment, T cells expressing Ortho-Receptor 2 specifically trafficked to the tumor transduced with Ortho-Ligand-encoding AAV (right-flank tumor) over a matched tumor transduced with a mock-payload virus (left-flank tumor) (**Fig. 5B, fig. S14**). These studies show that the Ortho-Ligand/Receptor 2 pair can be used to guide engineered T cell trafficking in vivo.

### Orthogonal chemokine system can spatially coordinate interaction between two cell populations in vivo

Immune responses often rely on coordinated interactions between multiple distinct cell types, which can only occur once these cells find each other within a complex in vivo environment (see Shah *et al*., companion paper, submitted).). Chemokines can normally guide this process by directing multiple cell populations toward the same spatial location (e.g. to lymph nodes where two cell populations can interact). Here we tested whether Ortho-Ligand could co-localize two interacting T cell populations – an engineered sender and receiver pair – to coordinate their communication in vivo. Sender T cells presented a membrane-tethered ALFA epitope tag as a model signal, and receiver T cells expressed an anti-ALFA SNIPR (synthetic intramembrane proteolysis receptor) (*14*), a synNotch family receptor, to sense the ALFA tag. The receiver cells were engineered to express an mCherry or effLuc reporter only when they received the sender cell signal.

To first test if the Ortho-Ligand could drive the productive interaction of these Sender-Receiver cells in vitro, we made wells containing a tumor spheroid expressing the Ortho-Ligand. A mixture of sender and receiver T cells were then added to these wells. We systematically varied whether the sender and receiver cells expressed Ortho-Receptor 2. We found that the receiver cells were only successfully activated in the spheroid when both sender and receiver T cells expressed Ortho-Receptor 2 and thereby co-localized to the tumor spheroid (**Fig. 5C**).

We then tested the ability of the Ortho-Ligand to induce this specific cell-cell interaction in vivo. NSG mice were engrafted with subcutaneous A549 tumors on both flanks, one producing a mock payload (mCherry) and the other producing Ortho-Ligand, both in a doxycycline-inducible manner. One day after starting the doxycycline diet, mice received both sender and receiver T cells intravenously (tail vein). Only when both T cell populations expressed Ortho-Receptor 2 did receiver T cells detect sender T cells, turning on the effLuc reporter specifically in the Ortho-Ligand-producing tumor. Expression of effLuc was absent when either Ortho-Receptor 2 or Ortho-Ligand was not expressed, demonstrating that the Ortho-Ligand/Receptor 2 pair enables effective cell-cell communication by spatially co-localizing the two populations in vivo (**Fig. 5D, fig. S15**). Thus, the ortho-chemokine system can be used to drive two cell populations to the same location in vivo, thereby strongly favoring their interaction, akin to what happens at key immune organs like lymph nodes.

### Engineering cell “swarming”: Ortho-chemokine positive feedback circuit that autonomously amplifies T cell recruitment

One of the unique features of the ortho-chemokine system is that one can engineer new orthogonal circuits that manipulate which cells make and which cells sense the migratory signal. Swarming circuits underlie a class of spatially self-organizing behaviors characterized by agents that simultaneously produce and sense an attractive signal. Swarming is observed in insects such as ants, which can produce pheromones that, in turn, recruit more ants. Similar swarming behaviors are observed at the cellular level, as seen in diverse cell types such as Dictyostelium and neutrophils (*15–18*). In these cases, a collective of cells both produces and senses chemotactic signals, yielding a self-organized gradient through positive feedback (*19, 20*).

We asked whether we could engineer a synthetic T cell trafficking circuit that uses positive feedback, mediated by the ortho-chemokine system, to amplify T cell recruitment to a target tissue (**Fig. 6A**). Such a circuit could serve as the basis for improved recruitment of engineered T cells to a tumor or other disease tissue. At its core, a swarming circuit would have a T cell engineered to both sense Ortho-Ligand (i.e. by expressing Ortho-Receptor) and produce Ortho-Ligand, in an input-induced manner. Such a circuit would require three elements: 1) an initiating signal to recruit a small number of nucleating “pioneer” T cells to the target site; 2) a target-encoded signal that induces these T cells to produce the Ortho-Ligand signal; 3) Ortho-Receptor expression to sense the Ortho-Ligand signal and sustain targeted recruitment of engineered T cells, together yielding a positive feedback swarming circuit.

**Fig. 6.**
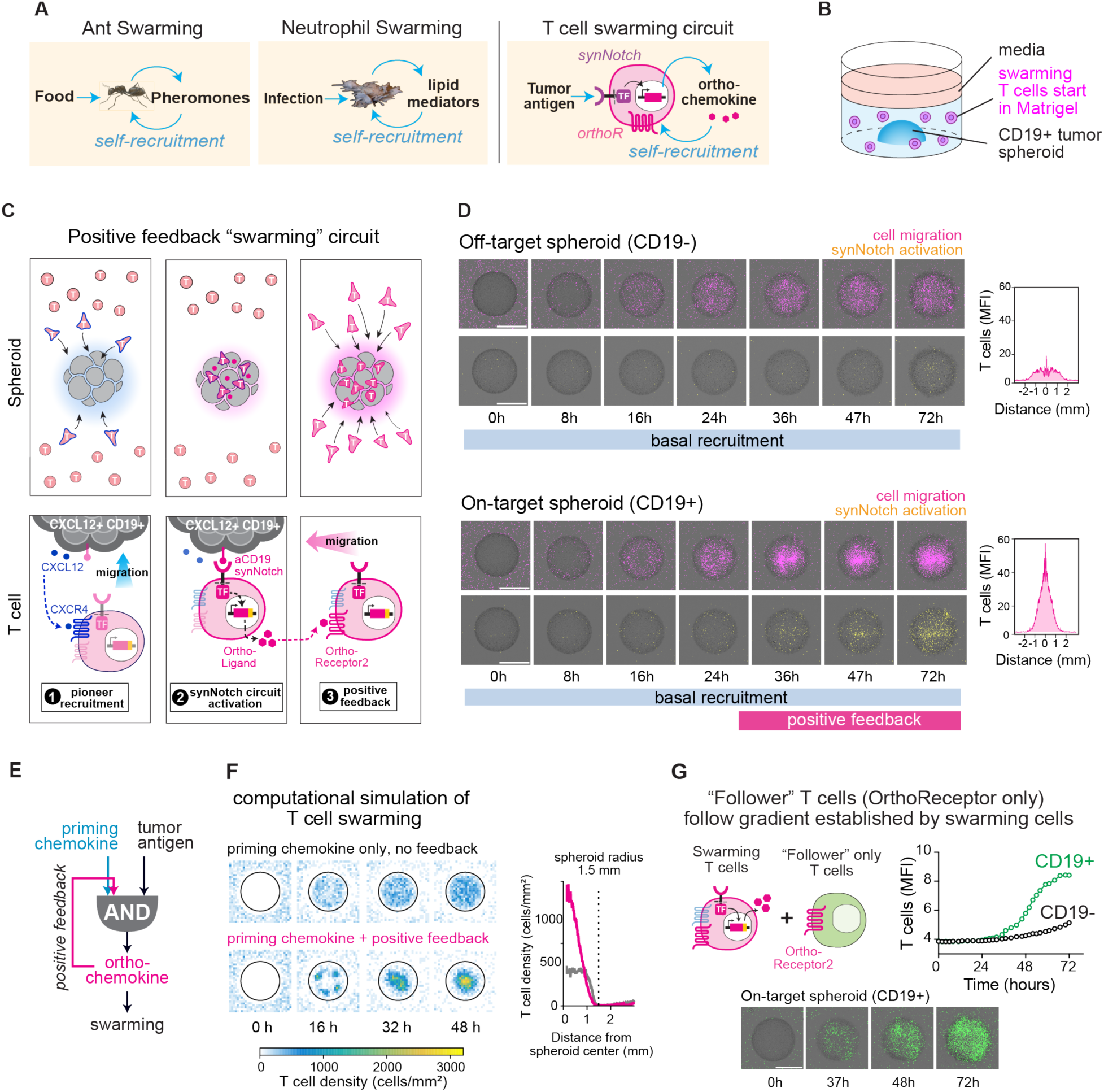
Ortho-Ligand/Receptor pair enables self-organizing collective behavior through a synthetic positive feedback “swarming” circuit. (**A**) Swarming in nature and by design. Ants and neutrophils self-recruit to target sites (e.g. food, infection) via positive feedback driven by pheromones and lipid mediators, respectively. This behavior can be synthetically reconstructed in T cells through positive feedback driven by synNotch-gated ortho-chemokine secretion and Ortho-Receptor-mediated sensing and migration. (**B**) T cell swarming is tested in a spheroid infiltration assay in a 24-well plate, with CXCL12-secreting A549 cells included at 10% of total cells to provide a priming signal. T cells are added in an overlying Matrigel and imaged over 72 hours. (**C**) Circuit schematic: (1) pioneer T cells are recruited via non-specific priming chemokine (CXCL12); (2) synNotch receptors detect tumor antigen (CD19) and induce Ortho-Ligand expression; (3) Ortho-Receptor 2 senses Ortho-Ligand and recruits additional engineered cells, establishing positive feedback that yields targeted, collective T cell “swarming.” (**D**) Timelapse images of T cell migration (magenta) and synNotch activation (yellow) over 72 hours. Scale bars, 2 mm. Spatial distribution of T cells within the spheroid, measured by the average fluorescence intensity line profile over 180° with 1° increments, shows T cells highly clustered at the CD19+ spheroid center. Representative of at least three independent experiments. See **fig. S16** for repeat data showing distinct T cell spatial distribution patterns. See **Movies S4, S5** for movies of T cell swarming. (**E**) AND-gate integration of priming chemokine and tumor antigen drives ortho-chemokine secretion, which feeds back as an input to the same AND gate, sustaining circuit activation and swarming. (**F**) Computational simulation of the swarming circuit recapitulates experimental results. T cell recruitment (N=10,000 cells) is simulated over 72 hours around a tumor spheroid (radius=1.5 mm). See **fig. S20, table S2,** and **Movie S8** for simulation details. (**G**) Follower T cells (green) expressing only Ortho-Receptor 2 (CXCR4^-^) migrate along the Ortho-Ligand gradient established by co-cultured swarming T cells (magenta; **fig. S18;** see **Movie S7**).

We could engineer this type of circuit using a synNotch receptor that recognizes a target tissue-specific antigen, and in response, induces expression of Ortho-Ligand. Specifically, T cells were engineered to constitutively express Ortho-Receptor 2 and an anti-CD19 synNotch receptor that induces expression of Ortho-Ligand upon target antigen binding (**fig. S10C**). A549 tumor spheroids were used as the target tissue (**Fig. 6B**) and were engineered to express membrane-tethered CD19 as a model antigen to drive synNotch-induced secretion of Ortho-Ligand. The A549 tumor spheroids were also engineered to secrete CXCL12 as an initiating attractive signal (**fig. S10D**), since T cells recognize this chemokine with the endogenously expressed CXCR4 receptor. We postulated that CXCL12/CXCR4 signaling would be able to non-specifically recruit an initial wave of T cells (“pioneer T cells”) to the tumor. Upon reaching the tumor and detecting CD19 through synNotch, pioneer T cells would then be induced to secrete Ortho-Ligand, establishing a self-generated Ortho-Ligand gradient specific to the target site. Because the T cells also express Ortho-Receptor 2, more of the engineered cells would migrate along this Ortho-Ligand gradient, and, in turn, produce additional Ortho-Ligand once they reached the tumor and detected CD19 antigen. This positive feedback loop could essentially convert the non-specific CXCL12-initiated trafficking into strongly amplified, antigen-specific (CD19+) T cell recruitment (**Fig. 6C**). Importantly, we would expect that a tumor spheroid lacking the synNotch antigen CD19 would show only low-level basal T cell recruitment, and no amplified swarming.

Indeed, timelapse imaging showed that engineered T cells bearing the synthetic circuit exhibited highly amplified swarming behavior towards a CD19-positive tumor spheroid (**Fig. 6D, fig. S16, Movies S4, S5**). The T cells showed a two-phase response, with weak initial recruitment, followed by highly amplified second phase of recruitment. This second phase coincided with activation of the synNotch receptor (detected via a mCherry fluorescence reporter co-induced by synNotch). In contrast, recruitment to the CD19-negative tumor spheroid showed only the initial weak recruitment phase (driven only by the basal priming chemokine CXCL12) and lacked the circuit-induced second wave of amplified cell recruitment. Thus, these cells were able to convert recognition of a tumor-specific antigen into a self-organized and amplified Ortho-Ligand gradient centered on the tumor (**Fig. 6E**).

Interestingly, with the Ortho-Ligand-driven swarming circuit, recruited T cells within the spheroid showed a highly focused spatial distribution (**Fig. 6D, fig. S16**). We computationally simulated the dynamics of this circuit (see **fig. S20, table S2** and **Materials and Methods** for details; see **Movie S8**). The simulation also showed the spatial pattern in which follower T cells nucleated around a few focal points where pioneer T cells were initially activated, before later merging into a single larger cluster around the spheroid center (**Fig. 6F**). This pattern was consistent with our experimental observation and can be explained by the positive feedback circuit acting to amplify both the number of recruited T cells, and the local production of ortho-chemokine in a non-linear, cell-density-dependent manner.

We also explored whether pioneer cell recruitment by a priming chemokine was necessary to initiate swarming. In the absence of a priming chemokine (CXCL12) in the tumor spheroids, swarming circuit activation was less reliable and seemed to rely on stochastic encounters with the CD19-positive spheroid. Without the priming chemokine, the swarming behavior was observed when we used a smaller well, where T cells had a higher probability of randomly entering the spheroid. In this situation, however, triggering of the positive feedback recruitment circuit occurred with delayed kinetics and a weaker magnitude (**fig. S19**). Thus, the priming chemokine is not absolutely necessary, but serves to increase the probability of recruiting pioneer cells to the target tissue, which can then set off the amplifying swarming circuit upon recognition of the synNotch target antigen.

### Swarming recruitment is selective for cells expressing Ortho-Receptor

Importantly, we found that T cells that did not express Ortho-Receptor 2 failed to be co-recruited as part of the second wave of amplified cell recruitment, even in the presence of intact swarming cells (**fig. S17, Movie S6**). This result shows that the amplified swarming recruitment is only selective for cells engineered to express the Ortho-Receptor 2. The cells that only recognize the priming chemokine but not the Ortho-chemokine (which would include endogenous immune cells that sense the priming chemokine) are not recruited as part of the amplified swarming phase.

Conversely, we found that if we added “follower” cells that only express Ortho-Receptor 2 (without the full swarming circuit), they were able to follow swarming cells and infiltrate the tumor spheroid as part of the amplified swarming phase. These follower cells (only expressing the Ortho-Receptor) migrated specifically to the CD19-positive spheroid during the second wave of amplified cell recruitment (**Fig. 6G, fig. S18, Movie S7**), but only when co-cultured with T cells with the complete swarming circuit. Follower cells also did not migrate to CD19-negative spheroids in the presence of swarming cells. Thus, following behavior is completely dependent on first establishing the amplified swarming gradient. These results indicate that once a swarming cell population has self-organized a Ortho-chemokine gradient, other follower cell species can be co-recruited to the target tissue. Thus, not all cells to be recruited need to express the full swarming circuit to be specifically trafficked to target sites.

## DISCUSSION

### New tools to precisely and selectively manipulate cell trafficking

We have engineered orthogonal chemokine-receptor pairs that can be used to rewire immune cell trafficking, with minimal crosstalk with the native chemokine network. This rewired specificity directly addresses a central limitation of the native chemokine network, its extensive redundancy and cross-interaction, which has confounded efforts to predictably control cell trafficking. Unlike native chemokines, Ortho-Ligand recruited only cells engineered with its cognate partner receptor, and did not recruit endogenous immune cells, even those recruited by its closest native relative, CCL20. We have demonstrated that T cells expressing the Ortho-Receptor can be specifically recruited to target tumors that express the Ortho-Ligand (e.g. via viral modification). This work thus introduces a new type of manipulation for controlling engineered immune cells – rather than modifying direct molecular recognition of target cells (e.g. CAR interaction with tumor antigen), these tools allow one to program the macroscopic behavior of where anatomically a cell population will traffic within the complex environment of the body, a critical feature of immune cell function and targeting specificity. Importantly, other non-imune cells also utilize chemokine trafficking, especially during development and regeneration. Thus, this type of orthogonal chemokine system could be used to control the trafficking of a broader set of immune and non-immune cells capable of migration.

### Engineering cell migration circuits capable of autonomous spatial self-organization

This engineered molecular pair represents one of the first orthogonal GPCR/ligand pairs, where both ligand and receptor are genetically encoded. This feature allows one to manipulate both what cells produce the signal and what cells detect it. There has been a long history of engineering GPCRs activated by orthogonal small molecules (*21–24*) or by light (*25*). In these cases, however, only the receptor is genetically encoded in target cells, and the input (drug/light) must be externally provided by the hand of a user. This feature has two limitations. First, it is difficult to use such systems to manipulate cells in vivo, as slow-release beads or light are constrained by factors such as surgical accessibility, tissue penetration, and the need for continual reapplication. Genetically encoded chemokines, in contrast, allow sustained, cell-autonomous production from spatially defined sender cells, in the simplest case, through established delivery platforms such as AAV or oncolytic viruses. Second, with genetically encoded receptors alone, it is not possible to engineer higher-order collective circuits that use feedback and other mechanisms to carry out autonomous, self-organizing behaviors. As demonstrated here, with genetically encoded ligand and receptor, we are able to engineer a synthetic swarming circuit, in which a T cell that induces ortho-chemokine via a synNotch receptor can promote its own positive feedback recruitment via the corresponding ortho-receptor, without the need for any user intervention. Thus, these types of genetically encoded orthogonal GPCR/ligand pairs provide a powerful new class of tools not only to modulate immune cell trafficking, but to engineer broader classes of autonomous GPCR-mediated cell-cell communication programs (**Fig. 7**).

**Fig. 7.**
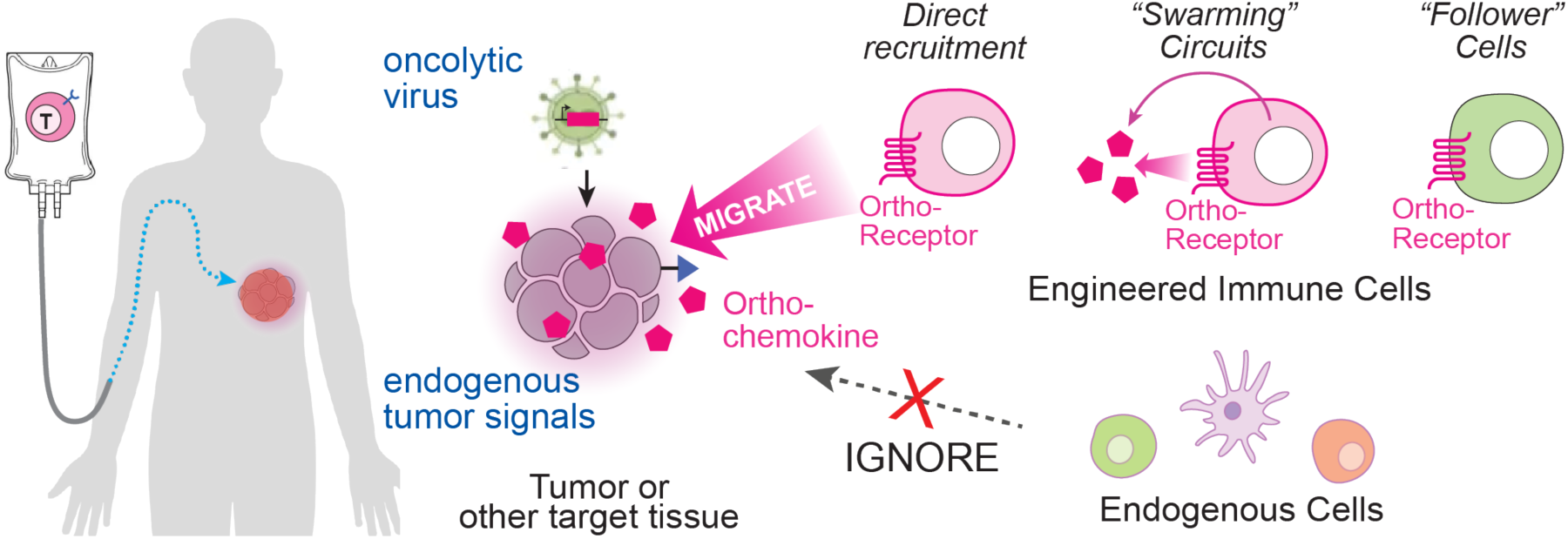
Diverse ways to utilize ortho-chemokine components to direct engineered immune cell trafficking. Local ortho-chemokine gradient can be established by various approaches, such as oncolytic virus gene delivery and cell-based protein delivery (e.g. synNotch circuit T cells). Only the engineered cells that express ortho-receptor can migrate along this gradient, and may further amplify targeted, collective migration through a positive feedback “swarming” circuit.

Here we take advantage of this orthogonal ligand/receptor pair to show how we can design synthetic circuits in which cells self-generate their own chemokine gradients, dynamically shaping their trafficking response as a collective. This work shows how these new tools enable exploration of more complex migration behaviors, beyond those observed with a fixed, externally imposed ligand gradient. Collective migration programs like these have been described in many native biological systems, including zebrafish primordial germ cell development, cancer cell metastasis, and immune cell trafficking (*15–20, 26–29*). In these systems, gradients are amplified through positive feedback in which migrating cells secrete additional chemokine (*19, 20*), or sharpened by ACKRs that locally deplete chemokine (*12, 13*). Our engineered positive feedback circuit demonstrates that this type of logic can be implemented synthetically. Pioneer T cells recognizing a target antigen convert an initial non-specific trafficking response into a strong, antigen-targeted swarming response, analogous to collective behaviors observed in ant foraging and neutrophil swarming (*15, 17*). This amplified trafficking response is highly specific for engineered cells expressing ortho-receptors, leaving other undesired cells blind to ortho-chemokine gradients. This circuit shows how key target information, such as a priming chemokine signal and a tumor-specific antigen, can be leveraged to drive self-organization of an ortho-chemokine guidance gradient.

These new tools will enable the exploration and optimization of these and other cell trafficking programs, which could more effectively and precisely target therapeutic cells to sites of disease. More broadly, these tools provide a foundation for engineering synthetic multicellular systems in which distinct cell populations must spatially organize to communicate and coordinate collective functional responses (Shah *et al*., companion paper, submitted).

## Acknowledgments

We thank G.M. Allen, H.K. Bhargava, P. Abi Kheirs, Y. Wang, M.S. Simic, R.R. Schlimgen, M.J. Wedemeyer, N. Blizzard, M.P. Lopez, S. Sidhu, A. DeGuzman for assistance and advice. We thank Htet Khant at the USC Cryo-EM facility for data collection.

## Funding

This work was supported by the following.

American Cancer Society Postdoctoral Fellowship – Marianne B. Kipper Research Fund (JL)

National Cancer Center Postdoctoral Fellowship (JL)

Anthropic’s AI for Science Program – Claude Science Cohort (DG)

Achievement Rewards for College Scientists (ARCS) Foundation Fellowship (DHS) Shurl and Kay Curci Foundation PhD Scholarship (LOH)

NSF Graduate Research Fellowship Program (SW)

Luxembourg Institute of Health (LIH NanoLux Platform) (MS, AC)

Cancer Foundation Luxembourg (MS, AC)

Luxembourg National Research Fund FNR 20/15084569 (MS, AC)

Luxembourg National Research Fund FNR C23/BM/18068832 (MS, AC)

Luxembourg National Research Fund FNR C25/BM/19563724 (MS, AC)

Luxembourg National Research Fund FNR 25/19352183 (MS, AC)

NIH R37 AI058072 (BFV) NIH U01CA265697 (WAL)

## Author contributions

Conceptualization: JL, BFV, AM, WAL

Methodology: JL, KS, CLDT, DG, FCP, MS, AC, BFV, AM, WAL

Investigation: JL, KS, CLDT, DG, DHS, LOH, SW, MC, WY, YT, MS, AC, AM, WAL

Visualization: JL, KS, CLDT, DG, MS, AC, AM, WAL

Supervision: BFV, AM, WAL

Writing – original draft: JL, WAL

Writing – review & editing: JL, MS, AC, BFV, AM, WAL

## Data, code, and materials availability

All data are available in the main text or the supplementary materials. Reagents are available from the corresponding author upon reasonable request. Plasmids from this paper will be made available on Addgene. Code used for the simulations is available at https://github.com/DebrajGhose/2026-cell-swarming-synchemokine. Coordinates for the wildtype and orthogonal (Ortho 2) chemokine-receptor complexes have been deposited in the Protein Data Bank (PDB) under accession codes 38ZA and 38YZ, respectively.

## Supplementary Materials

### Materials and Methods

Figs. S1 to S20

Tables S1 to S9

References (*30–60*)

Movies S1 to S8

## References and Notes

1. A. Rot, U. H. von Andrian, Chemokines in innate and adaptive host defense: basic chemokinese grammar. Annual Review of Immunology 22, 891–928 (2004).

2. J. C. Cyster, Chemokines and cell migration in secondary lymphoid organs. Science 286, 2098–2102 (1999).

3. A. B. Kleist, M. Szpakowska, L. J. Talbot, G. Slodkowicz, D. Malinverni, M. A. Thomas, K. S. Crawford, D. J. McGrail, A. F. Dishman, M. J. Wedemeyer, M. Sluter, S. S. Yi, N. Sahni, F. C. Peterson, A. Chevigné, B. F. Volkman, M. M. Babu, Encoding and decoding selectivity and promiscuity in the human chemokine-GPCR interaction network. Cell 188, 3603–3622 (2025).

4. H. He, P. Yang, Y. Gao, Chemokines and chemokine receptors: the key regulators of tumor microenvironment. Immunity and Inflammation 2, 24 (2026).

5. M. Pan, X. Wei, X. Xiang, Y. Liu, Q. Zhou, W. Yang, Targeting CXCL9/10/11-CXCR3 axis: an important component of tumor-promoting and antitumor immunity. Clinical and Translational Oncology 25, 2306–2320 (2023).

6. N. Redjimi, C. Raffin, I. Raimbaud, P. Pignon, J. Matsuzaki, K. Odunsi, D. Valmori, M. Ayyoub, CXCR3+ T regulatory cells selectively accumulate in human ovarian carcinomas to limit type I immunity. Cancer Research 72, 4351–4360 (2012).

7. Z. Eshhar, T. Waks, G. Gross, D. G. Schindler, Specific activation and targeting of cytotoxic lymphocytes through chimeric single chains consisting of antibody-binding domains and the gamma or zeta subunits of the immunoglobulin and T-cell receptors. Proceedings of National Academy of Science U.S.A. 90, 720–724 (1993).

8. K. T. Roybal, L. J. Rupp, L. Morsut, W. J. Walker, K. A. McNally, J. S. Park, W. A. Lim, Precision tumor recognition by T cells with combinatorial antigen-sensing circuits. Cell 164, 770–779 (2016).

9. L. Morsut, K. T. Roybal, X. Xiong, R. M. Gordley, S. M. Coyle, M. Thomson, W. A. Lim, Engineering customized cell sensing and response behaviors using synthetic Notch receptors. Cell 164, 780–791 (2016).

10. D. J. Wasilko, Z. L. Johnson, M. Ammirati, Y. Che, M. C. Griffor, S. Han, H. Wu, Structural basis for chemokine receptor CCR6 activation by the endogenous protein ligand CCL20. Nature Communications 11, 3031 (2020).

11. J. Dauparas, I. Anishchenko, N. Bennett, H. Bai, R. J. Ragotte, L. F. Milles, B. I. M. Wicky, A. Courbet, R. J. de Haas, N. Bethel, P. J. Y. Leung, T. F. Huddy, S. Pellock, D. Tischer, F. Chan, B. Koepnick, H. Nguyen, A. Kang, B. Sankaran, A. K. Bera, N. P. King, D. Baker, Robust deep learning–based protein sequence design using ProteinMPNN. Science 378, 49– 56 (2022).

12. A. Chevigné, D. F. Legler, A. Rot, S. Sozzani, M. Szpakowska, M. Thelen, International Union of Basic and Clinical Pharmacology. CXVIII. Update on the nomenclature for atypical chemokine receptors, including ACKR5. Pharmacological Reviews 77: 100012 (2024).

13. M. Szpakowska, D. F. Legler, S. Offermanns, S. Sozzani, A. Rot, M. Thelen, A. Chevigné, Inclusion of ACKR5 in the systematic nomenclature of atypical chemokine receptors. Nature Reviews Immunology 25, 225–226 (2025).

14. I. Zhu, R. Liu, J. M. Garcia, A. Hyrenius-Wittsten, D. I. Piraner, J. Alavi, D. V. Israni, B. Liu, A. S. Khalil, K. T. Roybal, Modular design of synthetic receptors for programmed gene regulation in cell therapies. Cell 185, 1431–1443 (2022).

15. V. Chandra, A. Gal, D. J. C. Kronauer, Colony expansions underlie the evolution of army ant mass raiding. Proceedings of National Academy of Science U.S.A. 118, e2026534118 (2021).

16. S. Sawai, P. A. Thomason, E. C. Cox, An autoregulatory circuit for long-range self-organization in *Dictyostelium* cell populations. Nature 433, 323–326 (2005).

17. K. Kienle, T. Lämmermann, Neutrophil swarming: an essential process of the neutrophil tissue response. Immunological Reviews 273, 76–93 (2016).

18. K. M. Glaser, M. Mihlan, T. Lämmermann, Positive feedback amplification in swarming immune cell populations. Current Opinion in Cell Biology 72, 156–162 (2021).

19. J. L. Galeano-Nino, S. V. Pageon, S. S. Tay, F. Colakoglu, D. Kempe, J. Hywood, J. K. Mazalo, J. Cremasco, M. A. Govendir, L. F. Dagley, K. Hsu, S. Rizzetto, J. Zieba, G. Rice, V. Prior, G. M. O’Neill, R. J. Williams, D. R. Nisbet, B. Kramer, A. I. Webb, F. Luciani, M. N. Read, M. Biro, Cytotoxic T cells swarm by homotypic chemokine signalling. eLife 9, e56554 (2020).

20. J. Cremasco, S. S. Tay, G. Yang, A. Huber, D. Kempe, R. Tang, F. Colakoglu, P. L.H. Newman, X. Lin, L. D. Goldstein, C. J. Kearney, M. Biro, Natural killer cells swarm and cross-recruit cytotoxic T cells via CCR5. Cell Reports 45, 117415 (2026).

21. P. Coward, H. G. Wada, M. S. Falk, S. D. H. Chan, F. Meng, H. Akil, B. R. Conklin, Controlling signaling with a specifically designed Gi-coupled receptor. Proceedings of National Academy of Science U.S.A. 95, 352–357 (1998).

22. B. N. Armbruster, X. Li, M. H. Pausch, S. Herlitze, B. L. Roth, Evolving the lock to fit the key to create a family of G protein-coupled receptors potently activated by an inert ligand. Proceedings of National Academy of Science U.S.A. 104, 5163–5168 (2007).

23. J. S. Park, B. Rhau, A. Hermann, K. A. McNally, C. Zhou, D. Gong, O. D. Weiner, B. R. Conklin, J. Onuffer, W. A. Lim, Synthetic control of mammalian-cell motility by engineering chemotaxis to an orthogonal bioinert chemical signal. Proceedings of National Academy of Science U.S.A. 111, 5896–5901 (2014).

24. N. A. Kalogriopoulos, R. Tei, Y. Yan, P. M. Klein, M. Ravalin, B. Cai, I. Soltesz, Y. Li, A. Ting, Synthetic GPCRs for programmable sensing and control of cell behaviour. Nature 637, 230–239 (2025).

25. Y. Xu, Y.-M. Hyun, K. Lim, H. Lee, R. J. Cummings, S. A. Gerber, S. Bae, T. Y. Cho, E. M. Lord, M. Kim, Optogenetic control of chemokine receptor signal and T-cell migration. Proceedings of National Academy of Science U.S.A. 111, 6371–6376 (2014).

26. D. D. Nogare, K. Somers, S. Rao, M. Matsuda, M. Reichman-Fried, E. Raz, A. B. Chitnis, Leading and trailing cells cooperate in collective migration of the zebrafish posterior lateral line primordium. Development 141, 3188–3196 (2014).

27. B. Boldajipour, H. Mahabaleshwar, E. Kardash, M. Reichman-Fried, H. Blaser, S. Minina, D. Wilson, Q. Xu, E. Raz, Control of chemokine-guided cell migration by ligand sequestration. Cell 132, 463–473 (2008).

28. L. Tweedy, P. A. Thomason, P. I. Paschke, K. Martin, L. M. Machesky, M. Zagnoni, R. H. Insall, Seeing around corners: Cells solve mazes and respond at a distance using attractant breakdown. Science 369, 1075 (2020).

29. R. H. Insall, P. Paschke, L. Tweedy, Steering yourself by the bootstraps: how cells create their own gradients for chemotaxis. Trends in Cell Biology 32, 585–596 (2022).

30. Q. Wan, N. Okashah, A. Inoue, R. Nehmé, B. Carpenter, C. G. Tate, N. A. Lambert, Mini G protein probes for active G protein– coupled receptors (GPCRs) in live cells. Journal of Biological Chemistry 293, 7466–7473 (2018).

31. R. Luís, G. D’Uonnolo, C. B. Palmer, M. Meyrath, T. Uchański, M. Wantz, B. Rogister, B. Janji, A. Chevigné, M. Szpakowska, Nanoluciferase-based methods to monitor activation, modulation and trafficking of atypical chemokine receptors. Methods in Cell Biology 169, 279–294 (2022).

32. M. Meyrath, M. Szpakowska, J. Zeiner, L. Massotte, M. P. Merz, T. Benkel, K. Simon, J. Ohnmacht, J. D. Turner, R. Krüger, V. Seutin, M. Ollert, E. Kostenis, A. Chevigné, The atypical chemokine receptor ACKR3/CXCR7 is a broad-spectrum scavenger for opioid peptides. Nature Communications 11, 3033 (2020).

33. C. B. Palmer, L. Rospape, M. Meyrath, C. Crudden, M. Counson, A.-K. Rohlfing, L. Di Niro, A. A. Bartolome, C. Pinheiro, V. Klapp, E. Cassano, S. A. Laporte, S. J. Hill, J. Drube, C. Hoffmann, R. Leurs, M. Bouvier, M. P. Gawaz, A. Hendrix, E. Moussay, M. J. Smit, J. Paggetti, M. Szpakowska, A. Chevigné, Chemokine and opioid peptide scavenging through constitutive and ligand-induced release of ACKR3-bearing extracellular vesicles. Nature Communications (in press) (2026). Preprint available at bioRxiv: 10.64898/2026.06.18.733119.

34. J. Schindelin, I. Arganda-Carreras, E. Frise, V. Kaynig, M. Longair, T. Pietzsch, S. Preibisch, C. Rueden, S. Saalfeld, B. Schmid, J.-Y. Tinevez, D. J. White, V. Hartenstein, K. Eliceiri, P. Tomancak, A. Cardona, Fiji: an open-source platform for biological-image analysis. Nature Methods 9, 676–682 (2012).

35. J.-Y. Tinevez, N. Perry, J. Schindelin, G. M. Hoopes, G. D. Reynolds, E. Laplantine, S. Y. Bednarek, S. L. Shorte, K. W. Eliceiri, TrackMate: An open and extensible platform for single-particle tracking. Methods 115, 80–90 (2017).

36. J. Jumper, R. Evans, A. Pritzel, T. Green, M. Figurnov, O. Ronneberger, K. Tunyasuvunakool, R. Bates, A. Žídek, A. Potapenko, A. Bridgland, C. Meyer, S. A. A. Kohl, A. J. Ballard, A. Cowie, B. Romera-Paredes, S. Nikolov, R. Jain, J. Adler, T. Back, S. Petersen, D. Reiman, E. Clancy, M. Zielinski, M. Steinegger, M. Pacholska, T. Berghammer, S. Bodenstein, D. Silver, O. Vinyals, A. W. Senior, K. Kavukcuoglu, P. Kohli, D. Hassabis, Highly accurate protein structure prediction with AlphaFold. Nature 596, 583–589 (2021).

37. M. Mirdita, K. Schütze, Y. Moriwaki, L. Heo, S. Ovchinnikov, M. Steinegger, ColabFold: making protein folding accessible to all. Nature Methods 19, 679–682 (2022).

38. E. F. Pettersen, T. D. Goddard, C. C. Huang, E. C. Meng, G. S. Couch, T. I. Croll, J. H. Morris, T. E. Ferrin, UCSF ChimeraX: Structure visualization for researchers, educators, and developers. Protein Science 30, 70–82 (2021).

39. D. P. Staus, L. M. Wingler, M. Choi, B. Pani, A. Manglik, A. C. Kruse, R. J. Lefkowitz, Sortase ligation enables homogeneous GPCR phosphorylation to reveal diversity in β-arrestin coupling. Proceedings of National Academy of Science U.S.A.115, 3834–3839 (2018).

40. S. Maeda, A. Koehl, H. Matile, H. Hu, D. Hilger, G. F. X. Schertler, A. Manglik, G. Skiniotis, R. J. P. Dawson, B. K. Kobilka, Development of an antibody fragment that stabilizes GPCR/G-protein complexes. Nature Communications 9, 3712 (2018).

41. CryoSPARC Team, K. Barber, H. Bridges, S. Dawood, K. Elder, N. Frasser, F. Hu, S. Liu, M. McLean, R. Narine, T. Ng, V. Peretroukhin, A. Punjani, H. Snyder, K. Stachowski, W. Tempel, S. Virani, R. Waldo, C. Wang, N. Wong, End-to-end automation of repeat-target cryo-EM structure determination in CryoSPARC. bioRxiv 2025.10.17.682689 [Preprint] (2025). 10.1101/2025.10.17.682689.

42. D. Liebschner, P. V. Afonine, M. L. Baker, G. Bunkóczi, V. B. Chen, T. I. Croll, B. Hintze, L.-W. Hung, S. Jain, A. J. McCoy, N. W. Moriarty, R. D. Oeffner, B. K. Poon, M. G. Prisant, R. J. Read, J. S. Richardson, D. C. Richardson, M. D. Sammito, O. V. Sobolev, D. H. Stockwell, T. C. Terwilliger, A. G. Urzhumtsev, L. L. Videau, C. J. Williams, P. D. Adams, Macromolecular structure determination using X-rays, neutrons and electrons: recent developments in Phenix. Structural Biology 75, 861–877 (2019).

43. P. Emsley, B. Lohkamp, W. G. Scott, K. Cowtan, Features and development of Coot. Structural Biology 66, 486–501 (2010).

44. C. R. Søndergaard, M. H. M. Olsson, M. Rostkowski, J. H. Jensen, Improved Treatment of Ligands and Coupling Effects in Empirical Calculation and Rationalization of pKa Values. Journal of Chemical Theory and Computation 7, 2284–2295 (2011).

45. T. J. Dolinsky, J. E. Nielsen, J. A. McCammon, N. A. Baker, PDB2PQR: an automated pipeline for the setup of Poisson–Boltzmann electrostatics calculations. Nucleic Acids Research 32, W665–W667 (2004).

46. D. A. Case, H. M. Aktulga, K. Belfon, D. S. Cerutti, G. A. Cisneros, V. W. D. Cruzeiro, N. Forouzesh, T. J. Giese, A. W. Götz, H. Gohlke, S. Izadi, K. Kasavajhala, M. C. Kaymak, E. King, T. Kurtzman, T.-S. Lee, P. Li, J. Liu, T. Luchko, R. Luo, M. Manathunga, M. R. Machado, H. M. Nguyen, K. A. O’Hearn, A. V. Onufriev, F. Pan, S. Pantano, R. Qi, A. Rahnamoun, A. Risheh, S. Schott-Verdugo, A. Shajan, J. Swails, J. Wang, H. Wei, X. Wu, Y. Wu, S. Zhang, S. Zhao, Q. Zhu, T. E. Cheatham III, D. R. Roe, A. Roitberg, C. Simmerling, D. M. York, M. C. Nagan, K. M. Merz Jr., AmberTools. Journal of Chemical Information and Modeling 63, 6183–6191 (2023).

47. S. Jo, T. Kim, V. G. Iyer, W. Im, CHARMM-GUI: A web-based graphical user interface for CHARMM. Journal of Computational Chemistry 29, 1859–1865 (2008).

48. E. L. Wu, X. Cheng, S. Jo, H. Rui, K. C. Song, E. M. Dávila-Contreras, Y. Qi, J. Lee, V. Monje-Galvan, R. M. Venable, J. B. Klauda, W. Im, CHARMM-GUI Membrane Builder toward realistic biological membrane simulations. Journal of Computational Chemistry 35, 1997–2004 (2014).

49. J. Lee, X. Cheng, J. M. Swails, M. S. Yeom, P. K. Eastman, J. A. Lemkul, S. Wei, J. Buckner, J. C. Jeong, Y. Qi, S. Jo, V. S. Pande, D. A. Case, C. L. Brooks III, A. D. MacKerell Jr., J. B. Klauda, W. Im, CHARMM-GUI Input Generator for NAMD, GROMACS, AMBER, OpenMM, and CHARMM/OpenMM Simulations Using the CHARMM36 Additive Force Field. Journal of Chemical Theory and Computation 12, 405– 413 (2016).

50. M. J. Abraham, T. Murtola, R. Schulz, S. Páll, J. C. Smith, B. Hess, E. Lindahl, GROMACS: High performance molecular simulations through multi-level parallelism from laptops to supercomputers. SoftwareX 1**–**2, 19–25 (2015).

51. J. Huang, S. Rauscher, G. Nawrocki, T. Ran, M. Feig, B. L. de Groot, H. Grubmüller, A. D. MacKerell Jr., CHARMM36m: an improved force field for folded and intrinsically disordered proteins. Nature Methods 14, 71–72 (2017).

52. J. B. Klauda, R. M. Venable, J. A. Freites, J. W. O’Connor, D. J. Tobias, C. Mondragon-Ramirez, I. Vorobyov, A. D. MacKerell Jr., R. W. Pastor, Update of the CHARMM All-Atom Additive Force Field for Lipids: Validation on Six Lipid Types. Journal of Physical Chemistry B 114, 7830–7843 (2010).

53. W. L. Jorgensen, J. Chandrasekhar, J. D. Madura, R. W. Impey, M. L. Klein, Comparison of simple potential functions for simulating liquid water. Journal of Chemical Physics 79, 926– 935 (1983).

54. N. Michaud-Agrawal, E. J. Denning, T. B. Woolf, O. Beckstein, MDAnalysis: A toolkit for the analysis of molecular dynamics simulations. Journal of Computational Chemistry 32, 2319–2327 (2011).

55. T. Takekoshi, J. J. Ziarek, B. F. Volkman, S. T. Hwang, A Locked, Dimeric CXCL12 Variant Effectively Inhibits Pulmonary Metastasis of CXCR4-Expressing Melanoma Cells Due to Enhanced Serum Stability. Molecular Cancer Therapeutics 11, 2516–2525 (2012).

56. C. Klemann, L. Wagner, M. Stephan, S. von Hörsten, Cut to the Chase: A Review of CD26/Dipeptidyl Peptidase-4’s (DPP4) Entanglement in the Immune System. Clinical and Experimental Immunology 185, 1–21 (2016).

57. H. Miller, J. Cosgrove, A. J. M. Wollman, E. Taylor, Z. Zhou, P. J. O’Toole, M. C. Coles, M. C. Leake, High-Speed Single-Molecule Tracking of CXCL13 in the B-Follicle. Frontiers in Immunology 9, 1073 (2018).

58. IUPAR/BPS Guide to PHARMACOLOGY, CXCR4 Receptor: Detailed View. Object ID 71 (https://www.guidetopharmacology.org/GRAC/ObjectDisplayForward?objectId=71).

59. M. J. Miller, S. H. Wei, M. D. Cahalan, I. Parker, Autonomous T Cell Trafficking Examined in Vivo with Intravital Two-Photon Microscopy. Proceedings of the National Academy of Sciences 100, 2604–2609 (2003).

60. R. E. Waugh, E. Lomakina, A. Amitrano, M. Kim, Activation Effects on the Physical Characteristics of T Lymphocytes. Frontiers in Bioengineering and Biotechnology 11, 1175570 (2023).

